# Deficiency of intellectual disability-related gene *Brpf1* reduced inhibitory neurotransmission and *Map2k7* expression in GABAergic interneurons

**DOI:** 10.1101/2021.01.11.426233

**Authors:** Jingli Cao, Weiwei Xian, Maierdan Palihati, Yu Zhu, Guoxiang Wang, Yunli Xie, Guomin Zhou, Linya You

## Abstract

Intellectual disability is closely related to impaired GABA neurotransmission. *Brpf1* was specifically expressed in medial ganglionic eminence (MGE), a developmental niche of GABAergic interneurons, and patients with *BRPF1* mutations were mentally retarded. To test its role in development and function of MGE-derived GABAergic interneurons, we performed immunofluorescence staining, whole-cell patch-clamp, MGE transplantation and mRNA-Seq to understand its effect on neuronal differentiation, dendritic morphology, electrophysiology, migration and gene regulation, using mouse MGE-derived GABAergic interneurons infected with AAV-shBrpf1. We found a decreasing trend on parvalbumin^+^ interneuron differentiation. Moreover, increased firing threshold, decreased number of evoked APs, and a reduced amplitude of mIPSCs were observed before any significant change of MAP2^+^ dendritic morphology and *in vivo* migration appeared. Finally, mRNA-Seq analysis revealed that genes related to neurodevelopment and synaptic transmission such as *Map2k7* were dysregulated. Our results demonstrated a key role of *Brpf1* in inhibitory neurotransmission and related gene expression of GABAergic interneurons.

## Introduction

Intellectual disability (ID) usually refers to a series of neurodevelopmental disorders that are obviously impaired in intelligence and adaptive function due to brain hypoplasia or organic damage[1, 2], affecting approximately 2-3% of the total population[3], with a prevalence of up to 2% in children or adolescents[4, 5]. Intellectual disability not only causes a heavy burden on patients and families, but also brings a series of problems in public health, society and education. At present, studies have shown that there are around 1396 genes related to ID[6], and bromodomain and PHD finger-containing protein 1 (*BRPF1*) is one of them. In the last few years, a total of 40 clinical cases of mental retardation caused by *de novo* or inherited *BRPF1* mutations have been reported[7–11]. Yan et al. reported that 10 patients with *BRPF1* mutations had symptoms such as developmental delay, language expression disorder and mental retardation[7]. Another team found 6 patients with *BRPF1* mutations whose clinical manifestations were growth retardation, sagging eyelid and cerebellar malformations[8]. Recent studies have further confirmed that patients with *BRPF1* mutations often exhibited clinical symptoms such as ID, general developmental delay, gross motor delay, facial and ocular deformities (such as blepharoptosis and ptosis)[9,11].

*BRPF1* is a unique epigenetic regulatory factor, which contains two PHD fingers, one bromodomain and one PWWP domain. *BRPF1* can recognize different epigenetic markers and activate three histone acetyltransferases MOZ (also known as KAT6A), MORF (also known as KAT6B) and HBO1[12-14]. Our previous work indicated that mouse *Brpf1* plays a key role in early embryo development[15], forebrain development[16, 17], and embryonic hematopoietic stem cell maintenance[18]. Global knockout mice were lethal before E9.5 with defects such as neural tube closure, angiogenesis, and placental development[15]. Forebrain-specific loss of *Brpf1* led to early postnatal lethality and growth retardation[17], and it has also been found that loss of *Brpf1* can regulate neuronal migration, cell cycle progression and transcriptional control, resulting in abnormal hippocampal morphogenesis[16]. Other group reported that heterozygous loss of *Brpf1* led to reduced dendritic complexity in both hippocampal granular cells and cortical pyramidal neurons[19].

ID is closely related to the imbalance of inhibitory/excitatory neural circuits, especially the imbalance of inhibitory circuits[20-22]. GABAergic interneurons are the major cellular units of the inhibitory neural circuit[23, 24], and are mainly produced in the medial and caudal ganglionic eminence (MGE and CGE) and preoptic area (PoA) of the telencephalon[25, 26]. After differentiation, GABAergic interneurons migrate tangentially to the neocortex, with some of them to the cortical marginal zone[27]. MGE-derived GABAergic interneurons are mainly differentiated into somatostatin (SST)^+^ and parvalbumin (PV)^+^ interneurons[26, 28], and growth number reaches a peak at E13.5[27]. GABAergic interneurons stay on the MGE for a period of time, then mostly tangentially migrate at E15.5-E16.5. Our previous work showed that mouse *Brpf1* is specifically expressed in MGE[29]. Relatedly, the haploid mutation of *Brd1* (also called *Brpf2*) significantly reduced the number of PV^+^ interneurons in the anterior cingulate cortex of mouse[30].We hypothesized that *Brpf1* regulates MGE-derived GABAergic interneurons and affects inhibitory neural circuits, which will eventually lead to abnormal learning, memory and cognitive ability.

To test this, we applied mouse MGE-derived GABAergic interneurons and AAV-shBrpf1 to examine the effect of *Brpf1* deficiency on the differentiation, dendritic morphology, electrophysiological properties, migration, and gene expression of MGE-derived GABAergic interneurons by immunofluorescence staining, whole-cell patch-clamp, MGE cell transplantation and mRNA-seq. This study will help us understand the key role of *Brpf1* on the morphology and function of GABAergic interneurons, and provide us with new insights into the pathogenesis of ID caused by *BRPF1* mutations.

## Results

### Brpf1 deficiency led to a mild decreased trend on the differentiation of PV^+^ interneurons

To study the effect of *Brpf1* knockdown on the differentiation of MGE-derived GABAergic interneurons, we dissected mouse MGEs from E14.5 embryos for primary culture of GABAergic interneurons. Neurons were infected with AAV-scramble-GFP or AAV-shBrpf1-GFP at DIV3, and immunolabeling was performed at DIV14-15. We confirmed a knockdown efficiency of approximately 40% by RT-qPCR (Figure 1D). Then the proportion of GABA^+^GFP^+^, PV^+^GFP^+^ and SST^+^GFP^+^ neurons in GFP^+^ neurons was calculated (Figure 1). The results showed that the ratio of GABA^+^GFP^+^ neurons in GFP^+^ neurons reached almost 100%, indicating that the majority of the cultured MGE-derived neurons were GABAergic interneurons (Figure 1A and 1E). GABAergic interneurons are mainly differentiated into SST and PV interneurons[26, 28]. Thus, we examined the ratio of PV^+^ GFP^+^ or SST^+^GFP^+^ in total GFP^+^ neurons (Figure 1B-1C, 1F-1G), and found little effect on SST^+^ neuronal differentiation but a mild decreased trend on PV^+^ neuronal differentiation upon *Brpf1* knockdown.

**Figure 1.**
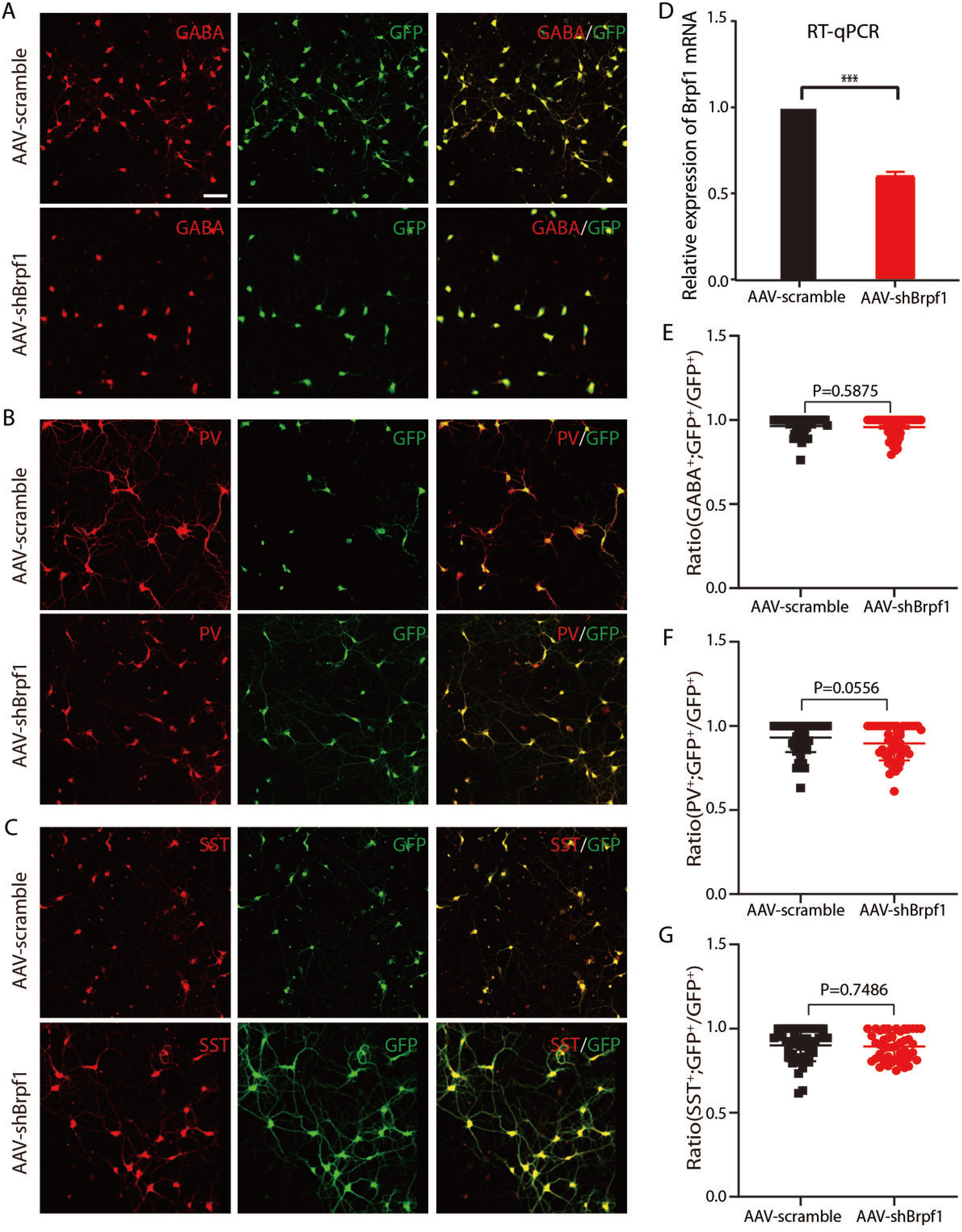
*Brpf1* knockdown led to a mild reduced trend on the differentiation of PV^+^ neurons from MGE-derived GABAergic interneurons. (A, B, C). Representative immunofluorescent images at DIV15, which were co-labeled with GABA and GFP, PV and GFP, SST and GFP antibodies, respectively. Scale bar, 100 μm. (D). Quantitative mRNA analysis showed that knockdown efficiency of *Brpf1* reached approximately 40% (AAV-scramble group, n = 5 batches; AAV-shBrpf1 group, n = 5 batches; *p*<0.001). (E, F, G). The ratio of GABA^+^GFP^+^, PV^+^GFP^+^, SST^+^GFP^+^ neurons in total GFP^+^ neurons upon Brpf1 knockdown was quantified respectively. (E). n = 43 and 52 fields for AAV-scramble and -shBrpf1, respectively, *p*=0.5875; (F). n = 55 and 50 fields for AAV-scramble and -shBrpf1, respectively, *p*=0.0556; (G). n=49 and 45 fields for AAV-scramble and -shBrpf1, respectively, *p*=0.7486). n was the number of fields, and the average cell count per field was about 25 neurons.

**Figure 2.**
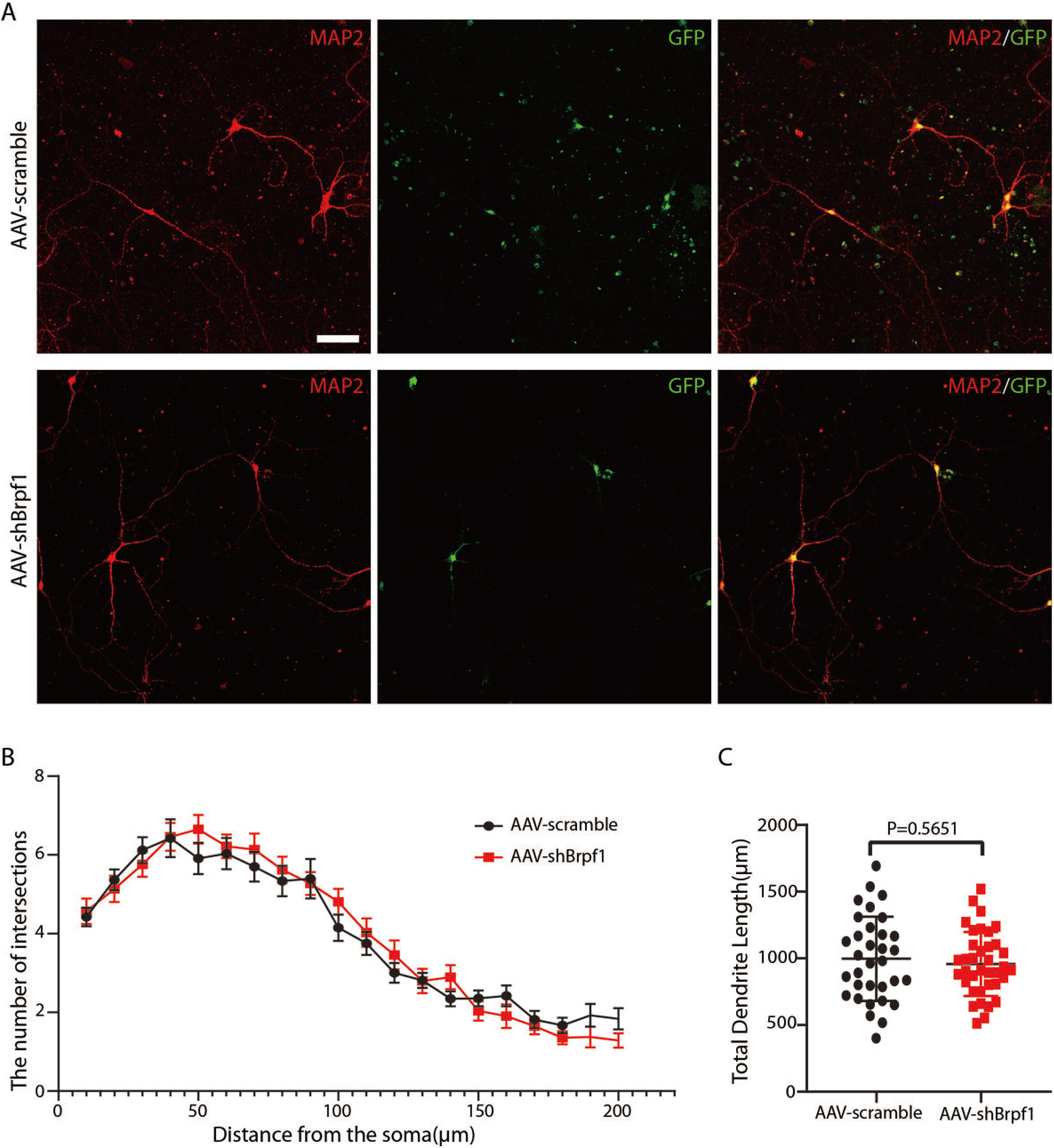
*Brpf1* knockdown had no significant effect on the dendritic morphology of MGE-derived GABAergic interneurons. (A). Representative immunofluorescent images of MGE-derived GABAergic interneurons immunostained with mouse anti-MAP2 and rabbit anti-GFP antibodies. Scale bar, 75 μm. (B). The mean number of intersections between dendrites and concentric circles with different radii (AAV-scramble group, n = 33 neurons; AAV-shBrpf1 group, n = 37 neurons). (C) Comparison of mean total length of neuronal dendrites (AAV-scramble group, n = 33 neurons; AAV-shBrpf1 group, n = 37 neurons; *p*=0.5651).

### Brpf1 deficiency had little effect on the dendritic morphology of MGE-derived GABAergic interneurons

The dendrites of neurons are essential for neuronal functions. Studies have shown that the complexity of neurites is closely related to some development-related diseases[31-33]. To study the effect of *Brpf1* knockdown on the dendritic morphology of MGE-derived GABAergic interneurons, we dissociated E14.5 MGEs and infected primary cultured neurons with AAV-scramble-GFP or AAV-shBrpf1-GFP. Immunostaining was performed at DIV 14-15 with anti-MAP2 antibody (a specific marker for dendrites) and the quantification was calculated from 3 independent experiments, totally 33 and 37 neurons in the AAV-Scramble and AAV-shBrpf1 group were analyzed, respectively, using Sholl analysis[34]. The results showed that there was no significant difference in the number of intersections and total length of dendrites between AAV-scramble and AAV-shBrpf1 groups, indicating that Brpf1 knockdown had little effect on the dendritic morphology of MGE-derived GABAergic interneurons.

### Brpf1 deficiency attenuated inhibitory neurotransmission by decreasing mIPSC amplitude and increasing firing threshold

Patients with ID usually have reduced learning and memory abilities, delay of which is closely related to synaptic dysfunction[35-37]. Synaptic dysfunction is usually determined by the characteristics of cell membranes and synaptic transmission between neurons. To assess the functional consequences of *Brpf1* knockdown on synaptic transmission, we performed whole-cell patch clamp to record the electrophysiological properties of MGE-derived GABAergic interneurons at DIV 15 after AAV-scramble or –shBrpf1 infection at DIV3 (Figure 3). For inhibitory neurotransmission, we measured miniature inhibitory postsynaptic current (mIPSC). Each neuron was recorded for 6 minutes and the signal of the later 3 minutes was selected for statistical analysis (Figure 3A, a short fragment of the signal was shown). The amplitude but not the frequency of mIPSC decreased significantly (Figure 3B-3C). For membrane characteristics, we measured resting membrane potential (RMP), action potential (AP), and evoked AP. RMP had little change (Figure 3D). Due to few or no spontaneous APs after many trials, we recorded evoked APs. The AAV-shBrpf1 group required a larger incident current to induce the evoked APs, with a significant increase in the firing threshold (Figure 3E), although the maximum frequency of evoked APs was not affected (Figure 3F). More specifically, when the evoked APs were induced against a serial step of increasing currents, there was a significant decrease in the number of spikes at current 130pA, 170pA and 190pA (Figure 3G). All the results indicated that *Brpf1* knockdown attenuated inhibitory neurotransmission of MGE-derived GABAergic interneurons.

**Figure 3.**
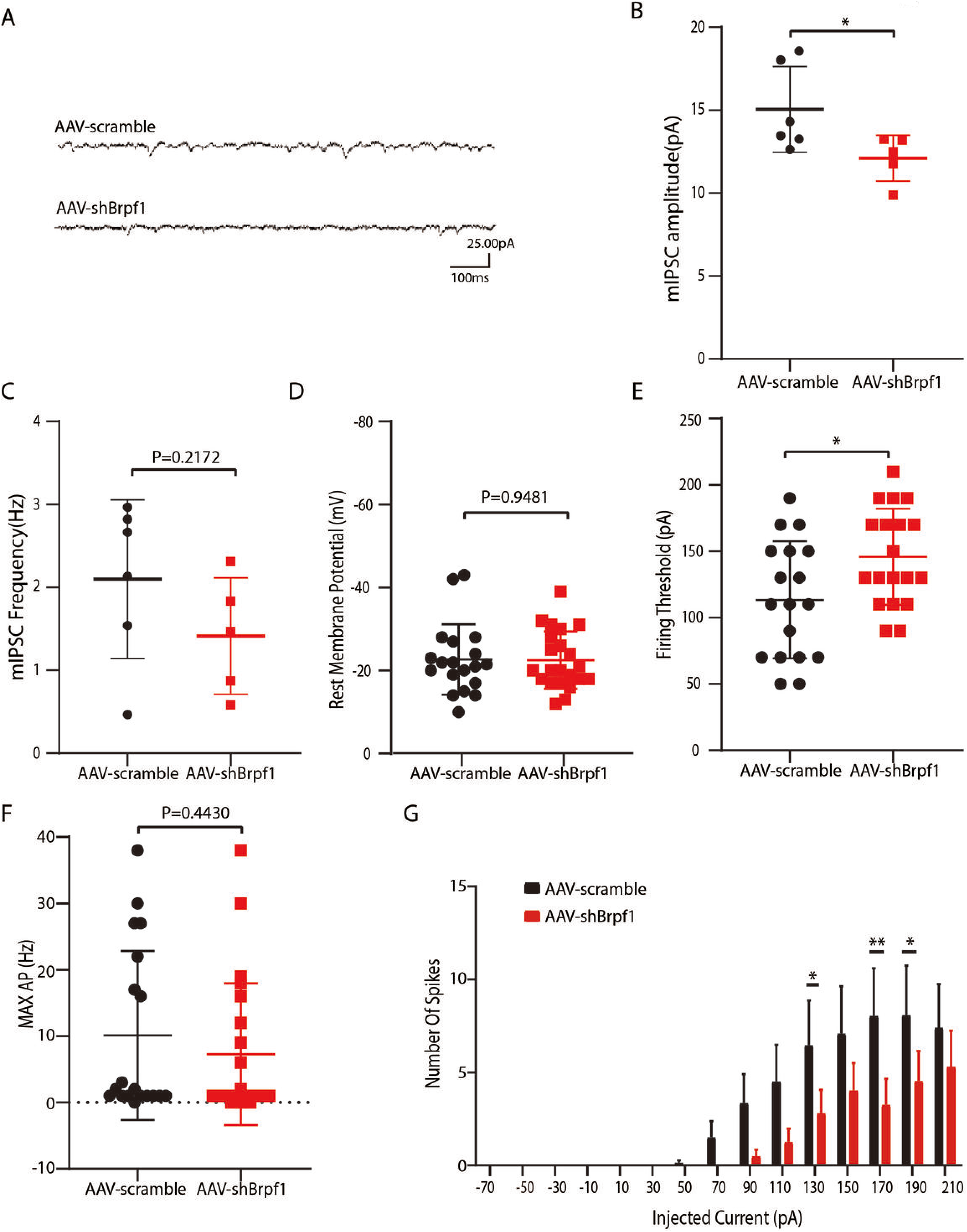
*Brpf1* deficiency led to attenuated inhibitory neurotransmission of MGE-derived GABAergic interneurons. (A) Representative traces of mIPSC in GABAergic interneurons at DIV 15 between the AAV-scramble and AAV-shBrpf1 groups. 25pA and 100ms were shown. (B) Statistical comparison of averaged mIPSC amplitude in GABAergic interneurons (AAV-scramble group, n = 6 neurons; AAV-shBrpf1 group, n = 5 neurons; *p*=0.0495).(C) Statistical comparison of averaged mIPSC frequency (AAV-scramble group, n = 6 neurons; AAV-shBrpf1 group, n = 5 neurons; *p*=0.2172). (D) Comparable resting membrane potentials were observed (AAV-scramble group, n = 19 neurons; AAV-shBrpf1 group, n = 22 neurons; *p*=0.9481). (E) The averaged firing threshold to induce evoked APs were compared (AAV-scramble group, n = 19 neurons; AAV-shBrpf1 group, n = 22 neurons; *p*=0.0193). (F) The maximum frequency of evoked APs showed little change (AAV-scramble group, n = 19 neurons; AAV-shBrpf1 group, n = 22 neurons; *p*=0.4430). (G) The number of evoked APs was induced against depolarizing current steps of increasing amplitude starting from −70 to 210 pA. (AAV-scramble group, n = 19 neurons; AAV-shBrpf1 group, n = 22 neurons; *p*= 0.03972 at 130pA; *p*= 0.0072 at 170pA; *p*= 0.0452 at 190pA).

### Brpf1 deficiency had little effect on GABAergic interneuron migration in vivo

Neurons after birth migrate to specific locations and form a network with surrounding cells to perform their specific functions. In order to study the localization and migration ability of MGE-derived GABAergic interneurons in the cortex upon *Brpf1* knockdown, we adopted a transplantation assay[38, 39]. Briefly, E13.5 MGE cells were infected with AAV-scramble-GFP or AAV-shBrpf1-GFP, then injected into the cortex of P1 wildtype mice by micro-syringe (Figure 4A). The transplanted cells were allowed to develop *in vivo* for 35 days (a time point at which mature interneuronal markers were expressed) and the proportion of GFP^+^ cells in the cortex was analyzed (Figure 4B-4D). First, we analyzed the deep (V and VI) and superficial (I-IV) layers of the cortex, and found the proportion of GFP^+^ cells showed no significant difference upon *Brpf1* knockdown (Figure 4C). Upon further dividing cortex layers into layer I, II-IV, V and VI, the ratio of GFP^+^ cells in each layer still had little change (Figure 4D). Thus, the mild knockdown of *Brpf1* did not affect the neuronal migration and cortical localization of MGE-derived GABAergic interneurons.

**Figure 4.**
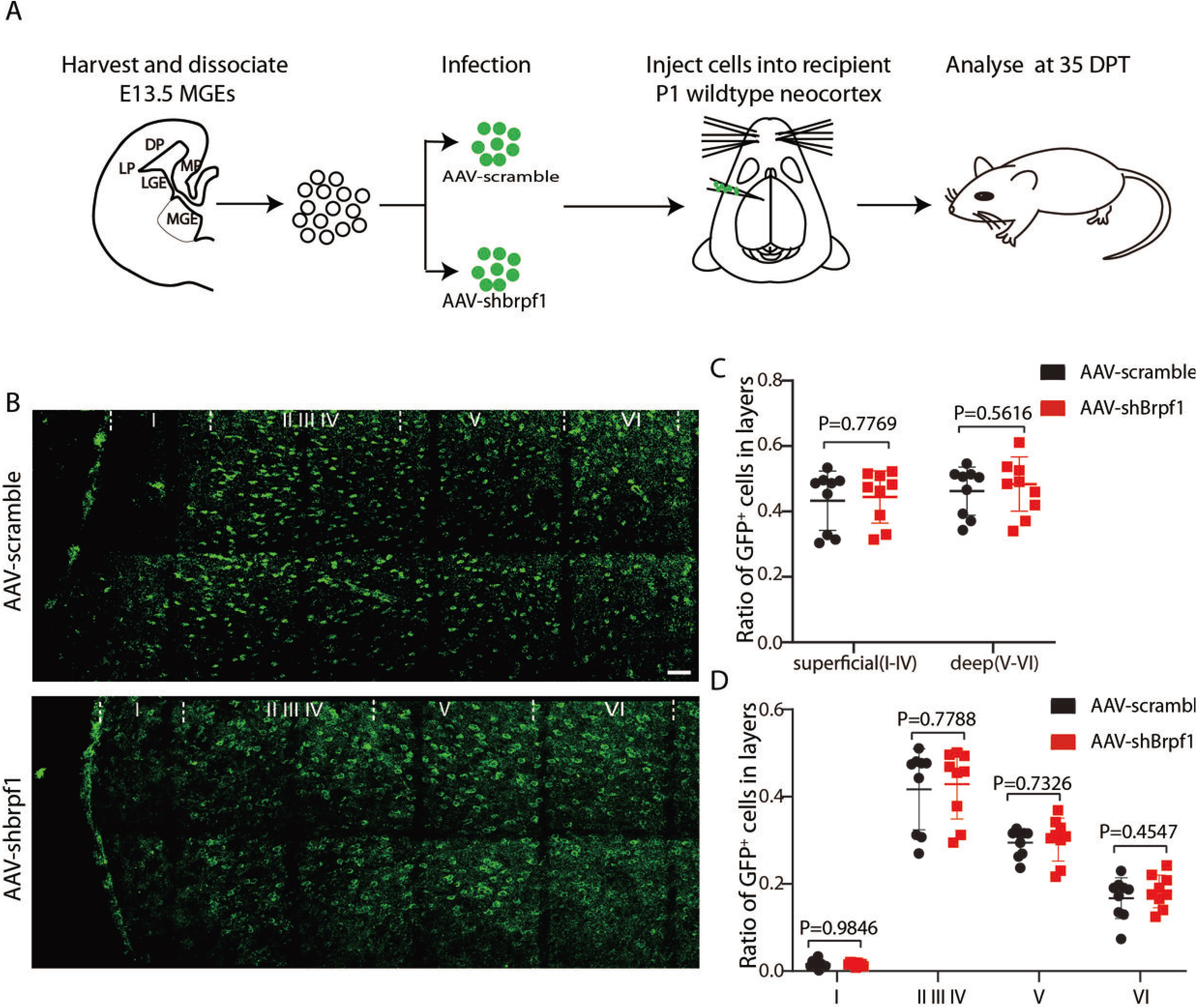
*Brpf1* deficiency had no effect on the neuronal migration of MGE-derived GABAergic interneurons *in vivo*. (A) Schematic diagram of MGE cell transplantation. In short, MGE neurons dissociated from E13.5 embryonic brains were infected with GFP-carrying adeno-associated virus (AAV-scramble-GFP or AAV-shBrpf1-GFP) and transplanted into the P1 neocortex. The proportion of GFP^+^ neurons in each cortical layer were analyzed after 35 days. (B) Representative images of GFP^+^ neurons in each layer of cortex analyzed at day 35 after transplantation. Arrows indicated GFP^+^ neurons. Scale bar: 50μm. (C) Quantitative analysis of the proportion of GFP^+^ neurons in deep (V and VI) and superficial (I-IV) cortical layer, respectively (AAV-scramble, n=9 mice; AAV-shBrpf1, n=9 mice). (D) Quantitative analysis of the ratio of GFP^+^ neurons in layer I, II-IV, V, and VI, respectively (AAV-scramble, n=9 mice; AAV-shBrpf1, n=9 mice).

### Brpf1 deficiency led to dysregulated MAPK pathway via Map2k7 in MGE-derived GABAergic interneurons

To study the molecular mechanism of *Brpf1* knockdown leading to changes in electrophysiological properties of MGE-derived GABAergic interneurons, three batches of RNA from AAV-scramble and AAV-shBrpf1 groups were extracted for mRNA-Seq. The number of fragments per kilobase fragment (FPKM) of the transcript per thousand fragments was used to calculate the expression of the known gene. DESeq2 software was used to screen the differentially expressed genes (DEGs) between different sample groups. DEGs with *p* < 0.05 and log_2_ (fold change) >=1 were considered to be up-regulated genes. Similarly, DEGs with *p* < 0.05 and log_2_ (fold change) <= −1 were considered to be down-regulated genes. 24 and 22 genes were upregulated and downregulated, respectively, upon Brpf1 knockdown (Figure 5A and Table S1).

**Figure 5.**
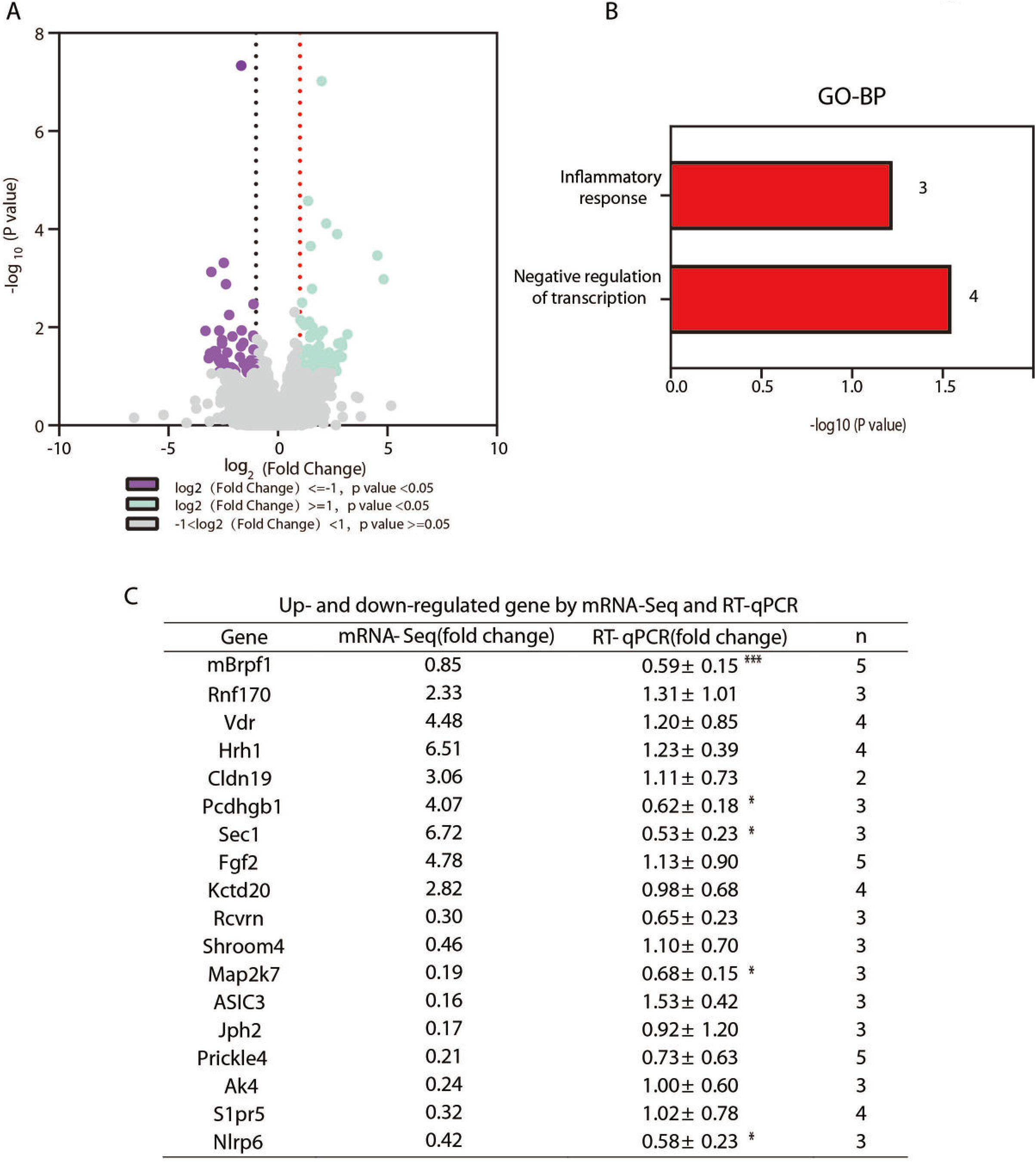
*Brpf1* deficiency led to decreased *Map2k7* expression in MGE-derived GABAergic interneurons revealed by mRNA-Seq. (A). Volcano plot of DEGs from mRNA-Seq, log_2_(fold change) <= −1 or >= 1 was set as significant downregulation or upregulation (AAV-scramble group, n = 3; AAV-shBrpf1 group, n = 3). (B). GO-analysis of 24 upregulated and 22 downregulated genes revealed by mRNA-Seq. The number of genes involved in each Go term was indicated. (C). Validation of selected DEGs from mRNA-Seq by RT-qPCR. n indicated pairs of scramble and shBrpf1 groups validated by RT-qPCR. *, *p*< 0.05, ***, *p*< 0.001.

GO and KEGG enrichment analysis of these genes revealed that most of the 24 up-regulated genes were involved in gene negative transcription and inflammatory response (Figure 5B and Table S1). Among them, *Tceal7*, *Mecom*, *Vdr*, *Lrch4* were involved in negative regulation of transcription, and *Hrh1*, *Mecom*, *Stab1* in inflammatory response. *Vdr* and *Hrh1* were verified by RT-qPCR (Figure 5C). However, for the 22 downregulated genes, no pathways were significantly enriched.

To further explore the underlying molecular mechanism, we selected most neuron-related genes among the 46 DEGs (24 plus 22) and validated by RT-qPCR (Figure 5C). For downregulated DEGs, *SHROOM4* was implicated in neural development, as its mutations were associated with human X-linked mental retardation[40]; ion transport related genes such as *Asic3*[41], *Ak4*[42], and *Slpr5*[43], can promote the release of cations to the outside of the cell, and thus a greater current stimulation is required to induce an AP; *Map2k7* is a key gene that activates MAPK signaling pathway[44], *Morf* (one of the three KATs *Brpf1* binds) can also specifically regulate this pathway[45]. Moreover, *Map2k7* heterozygous mice showed insufficient prefrontal cortex-dependent working memory[44, 46], consistent with the learning and memory impairment in Brpf1 mutant mice[19]; *Nlrp6* is part of *Nlrp6* inflammasome and plays a crucial role in innate immunity and inflammation. *Nlrp6* could induce apoptosis to regulate the survival of neurons[47].Interestingly, study found that the space exploration ability and associative learning of *Nlrp6*^−/−^ male mice were weakened[48]. We validated that *Map2k7* and *Nlrp6* were significantly decreased upon Brpf1 knockdown (Figure 5C). Of note, Both *Map2k7* heterozygous and tissue-specific knockout mice showed symptoms related to mental retardation, such as cognitive dysfunction and motor dysfunction (see discussion for details) [44, 49], which is similar as that of *Brpf1* knockout. For upregulated DEGs, overexpression of *Rnf170* inhibited IP3 receptor[50], which plays a key role in cell signaling; overexpression of transporter genes such as *Vdr[51]*, *Kctd20*[52], affects the assembly of some cation channels protein, and eventually leads to changes in membrane ion permeability.

## Discussion

Accumulating clinical studies has reported a total of 40 cases of patients with *BRPF1* monoallelic mutations, with symptoms such as mental retardation, developmental delay and epilepsy[7-11]. Impaired GABA neurotransmission is associated with mental disorders such as epilepsy and mental retardation. Our previous work has found that *Brpf1* global knockout is embryonically lethal[15], and *Brpf1* forebrain-specific knockout showed hypoplasia in hippocampus, corpus callosum and cortex[16, 17]. Moreover, we found that *Brpf1* is specifically expressed in MGE at E14.5[29], a niche where most GABAergic interneurons were born. To study monoallelic loss effect of *Brpf1*, we mildly knocked down *Brpf1* in MGE-derived GABAergic interneurons with the knockdown efficiency of about 40% (Figure 1D). Interestingly, this study found that *Brpf1* mild knockdown led to changes in electrophysiology (Figure 3) and PV^+^ neuronal differentiation (Figure 1) before any influence showed in dendritic morphology (Figure 2) and neuronal migration (Figure 4). More specifically, *Brpf1* mild knockdown can sensitively cause attenuated inhibitory neurotransmission by decreasing mIPSC amplitude and increasing firing threshold of evoked APs (Figure 3). The underlying mechanism may involve dysregulated MAPK pathway via *Map2k7*, as indicated by mRNA-Seq (Figure 5).

We found that the *p* value of the comparison of PV^+^ neurons is 0.056, suggesting a decreased trend of MGE-derived GABAergic interneurons to differentiate into PV^+^ interneurons upon *Brpf1* knockdown. Consistently, *Brd1* (also known as *Brpf2*) heterozygous mice also displayed a decrease in the number of PV^+^ interneurons[30, 53]. Moz and Morf (two of the three enzymes Brpf1 binds) were involved in regulating neurogenesis[54-56]. *Morf* mutant mice showed reduced ability of neural stem cells/progenitor cells differentiating into neurons, astrocytes, and oligodendrocytes, but not limited to specific neuronal types[54]. In our study, we observed a decreased trend of PV^+^ interneuronal differentiation upon mild *Brpf1* reduction. Further confirmation with a more efficient knockdown/knockout *in vivo* will be needed to study *Brpf1*’s effect on PV^+^ interneuronal differentiation.

Dysregulation of excitatory/inhibitory neuronal circuits was closely related to various neurodevelopmental disorders[57-59] such as Parkinson, intellectual disability, epilepsy, schizophrenia, etc. Our study found that insufficient *Brpf1* led to abnormal synaptic transmission in MGE-derived GABAergic interneurons with reduced amplitude of mIPSC and increased firing threshold of evoked AP. Consistently, previous studies showed that Brpf1 forebrain-specific (by Emx1-Cre) heterozygotes had increased firing threshold of AP, a downward trend of spike number, reduced mEPSC frequency and amplitude in hippocampus[19]. In addition, *Brd1*^+/-^ pyramidal neurons showed decreased frequency of spontaneous IPSCs and mIPSCs[60]. These results indicated that membrane properties and functional maturation of MGE-derived GABAergic interneurons were sensitive to *Brpf1* dose even with mild knockdown.

Interestingly, the mild knockdown dose of *Brpf1* in our study only led to significant changes in electrophysiology properties before any major influence showed in dendritic morphology and neuronal migration. Thus we performed mRNA-Seq analysis to mainly explain the change in electrophysiology. The analysis revealed that *Map2k7* and *Nlrp6* genes were significantly downregulated. Relatedly, *MOZ* knockdown led to a decrease in JNK(a key component of MAPK signaling pathway) phosphorylation[56], and patients with JNK mutations were detected with ID[61]. *MORF* disruption leads to a Noonan syndrome-like phenotype mainly via hyperactivated MAPK signaling in human and mice[45]. *Map2k7* also plays an important role in the regulation of MAPK signaling pathway[44]. In addition, *Map2k7* heterozygous mice exhibited cognitive dysfunction[44] and neuron-specific *Map2k7* knockout exhibited motor dysfunction such as muscle weakness and abnormal walking[49]. For *Nlrp6*, *Nlrp6*^−/−^ mice showed symptoms related to mental retardation[48]. The mechanism of how *Brpf1* regulates *Map2k7* expression merits further investigation.

This study helps us understand the key roles of *Brpf1* in differentiation, dendritic morphology, electrophysiological characteristics, migration and gene expression of MGE-derived GABAergic interneurons from a neurobiological level, and provides us with new insights into the pathogenesis of ID related to *BRPF1* mutations.

## Acknowledgements

We thank members from Dr. Linya You’s lab for their support and help during the experiment. This work was supported by the National Natural Science Foundation of China (81771228) to Dr. Linya You and National Natural Science Foundation of China (31571238) to Dr. Guomin Zhou.

## Materials and Methods

### Animals

C57BL/6 mice were used in this study and were purchased from Shanghai Slac Laboratory Animal CO., LTD. The day when vaginal plug is detected was considered to be embryonic day 0.5 (E0.5). All experimental animals were kept in an animal facility at Fudan University and all experiments were conducted in accordance with guidelines approved by Fudan University.

### Primary Neuron Culture, Immunofluorescence, and Quantification

Primary GABAergic interneurons were prepared from E14.5 wild-type pregnant mice. MGEs were isolated from the embryonic brains in ice-cold HBSS (Thermo,14185052) with 1% HEPES (Thermo, 15630106) and the MGEs were transferred to a 15 ml centrifuge tube for digestion with 3ml HBSS/HEPES containing 100 μg /ml DNase (Sigma, D4527) and 0.05% trypsin (Sigma, T8658-1vl). Centrifuge tube was transferred to a 37℃ water bath for 15min and gently shaked every 3-5 minutes. 5 ml Neurobasal medium (Gibco, 211030049) supplemented with 10% FBS (Gibco, 10091148), 2% B27 supplement (Gibco, 17504044), and 1% GlutaMAX-1(Gibco, 35050061) was added to stop the digestion. The mixed liquid was centrifuged at 800 rpm for 5 minutes and the supernatant was removed. Neurobasal medium supplemented with 10% FBS, 2% B27 supplement, and 1% GlutaMAX-1 was added again. The neurons were then dissociated by pipetting, and the suspension was centrifuged at 1000 rpm for 5 minutes. Cells were then resuspended, plated at a density of 2-5 × 10^4^ cells /well on coverslips in a 24-well plate pre-coated with poly-L-lysine (Sigma, P9155) and cultured in a Neurobasal medium supplemented with 2% B27 supplement, and 1% GlutaMAX-1.

Neurons were infected with AAV-scramble-GFP or AAV-shBrpf1-GFP virus at DIV3. The shBrpf1 sequence was shown in Table 1. After DIV10, GFP-expressing neurons could be gradually observed under a fluorescent microscope. At DIV14-15, immunofluorescent staining was performed to quantify the number of GABAergic interneurons that differentiated into PV^+^ or SST^+^ interneurons. Briefly, neurons were fixed in 4% paraformaldehyde (Biosharp, BL539A) in PBS (Hyclone, SH30243.01) for 20 minutes, permeabilized with 0.1% Triton X-100 (Sigma, X-100) in PBS for 1 hour, and blocked with 10% goat serum (Proteintech, b900780) in PBS for 1 hour at room temperature. Neurons were then incubated with primary antibodies at 4 ℃ overnight and second antibodies for 1 hour. The primary antibodies used were mouse anti-MAP2 (Proteintech, 67015-1-ig, 1:1000), rabbit anti-GFP (Proteintech, 50430-2-AP, 1:500), mouse anti-GABA (Sigma, A0310, 1:200), rabbit anti-PV (Proteintech, 26521-1-ap, 1:500), rabbit anti-SST (Proteintech, 17512-1-ap, 1:500), and mouse anti-GFP (Proteintech, 66002-1, 1:500). Neurons were finally incubated with DAPI (Sigma, D9542) at room temperature for 10 minutes and mounted with antifade mounting medium.

To study the effect of Brpf1 knockdown on the differentiation of GABAergic interneurons, we used ImageJ Fiji software to manually count the number of GABA^+^ GFP^+^, SST^+^GFP^+^, PV^+^GFP^+^ neurons and total number of GFP^+^ neurons, the proportion of co-labeled neurons in GFP^+^ neurons was then calculated.

**Table 1.**
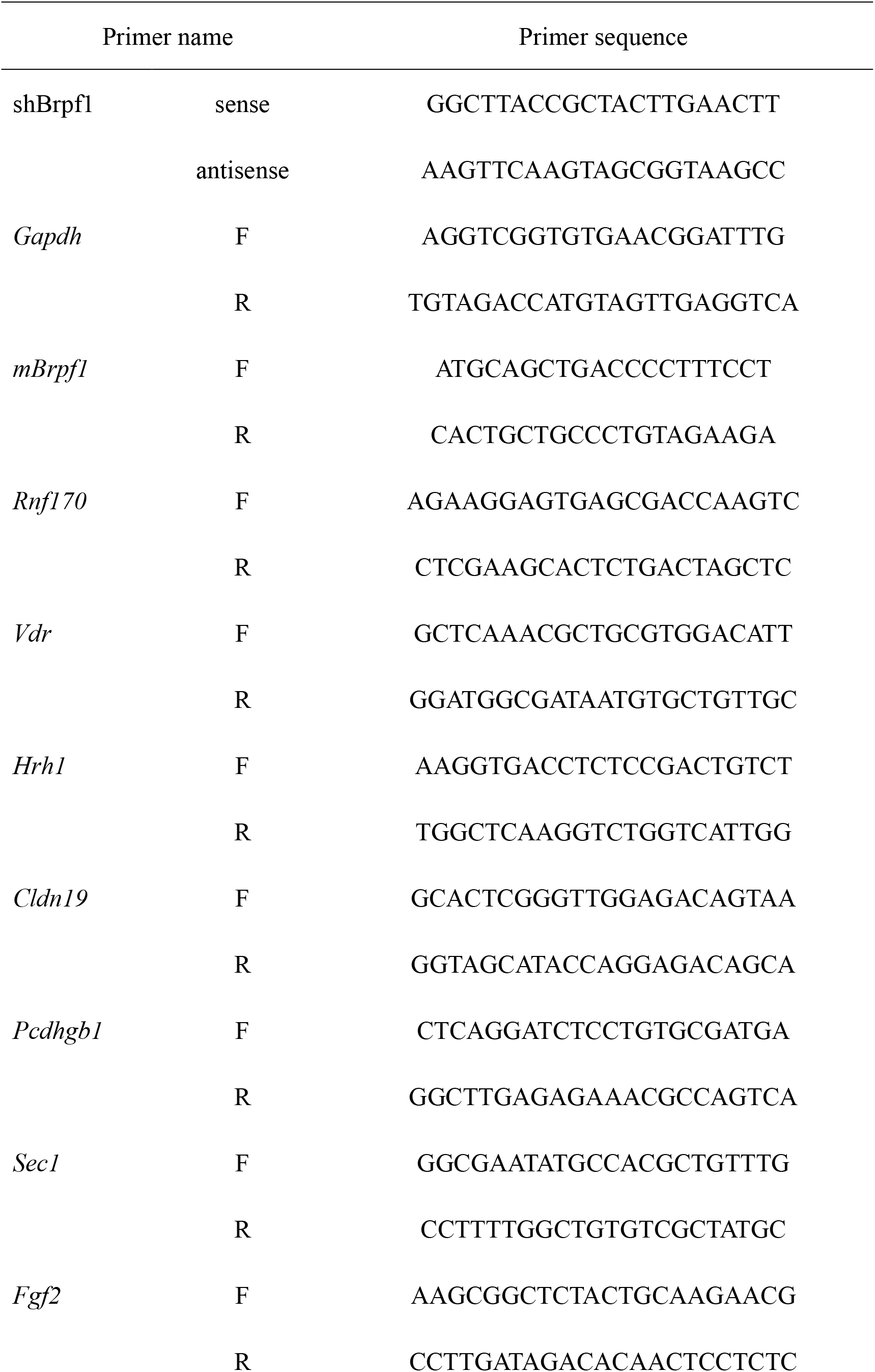
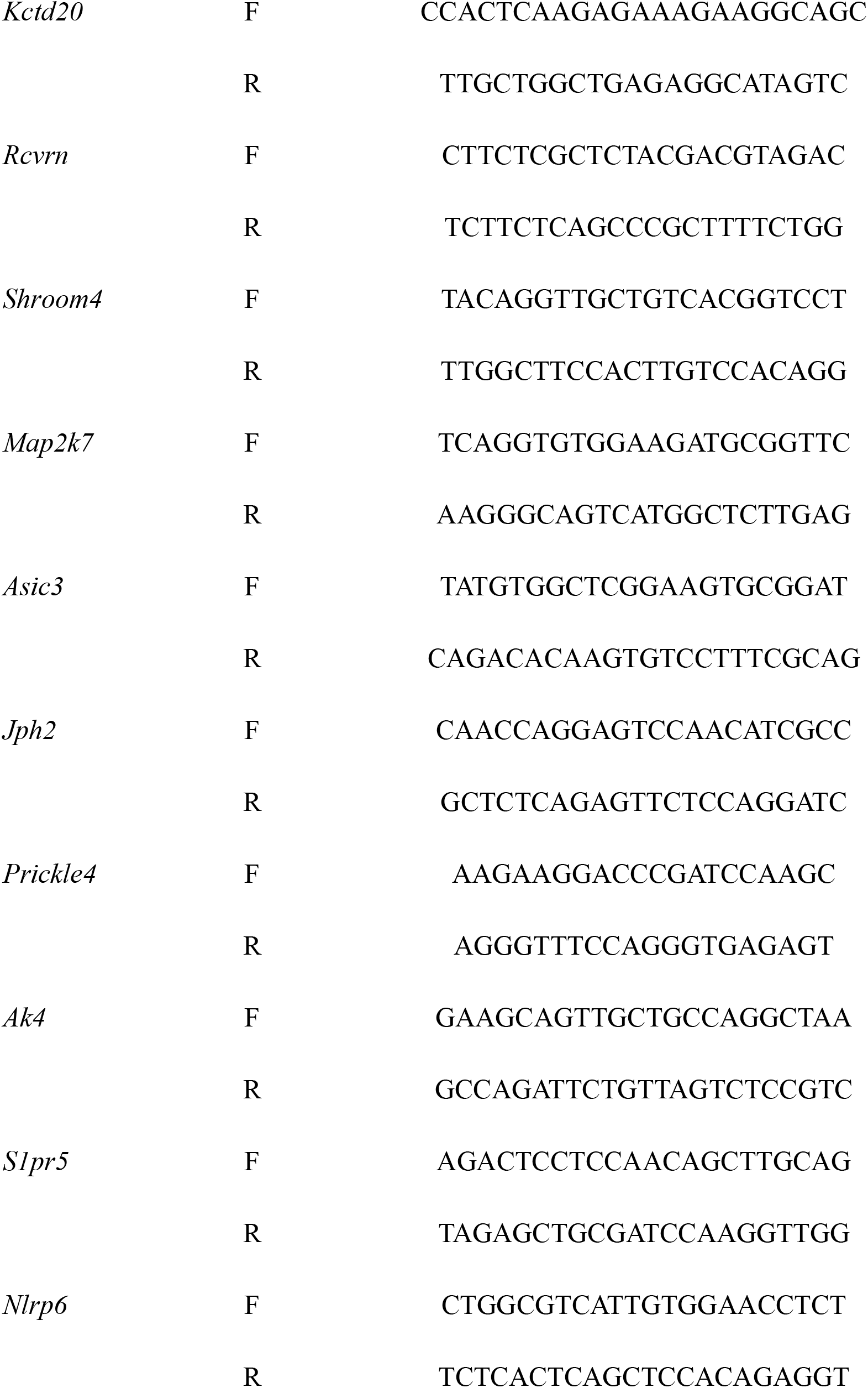
Primer sequence

For analysis of dendritic morphology, coverslips were observed under an inverted fluorescence microscope (Olympus) and confocal microscope (leica SP8). The images were analyzed using ImageJ Fiji. To calculate the branches and the total length of dendrites, Sholl analysis was applied. Simple Neurite Tracer plug-in was used to track the path of the neuron, and the radius was increased to cover the entire path of the neurons with the cell body as the center and a fixed value of 10um. The Sholl analysis plug-in was used to calculate the number of intersections between the ring and the neurite. The total length of dendrites was also displayed when tracing the path of neurites.

### Electrophysiological Recordings and analysis

For whole-cell patch-clamp recording, neurons were infected with a reduced titer of AAV-scramble-GFP or AAV-shBrpf1-GFP virus at DIV3 to achieve sparse labeling and the recordings were performed at DIV15. Neurons expressing GFP under high-resolution fluorescent microscope were chosen for electrophysiological recordings. Briefly, coverslips covered with neurons were transferred to a holding chamber attached to the microscope stage, containing normal-temperature artificial cerebrospinal fluid (ACSF). Electrodes with electrode solution were used to patch neurons under the fluorescent microscope. To characterize the intrinsic membrane properties of neurons, current clamp recordings were performed and hyperpolarized and depolarized current steps were injected in 20 pA increments. The following parameters were measured to characterize the neuronal membrane characteristics: the resting membrane potential was recorded immediately after the neuronal membrane was ruptured, and the number of APs induced after different current injections was measured. The AP current threshold is defined as the first rectangular current injection that causes a spike. Offline analysis was performed using Clampfit software. To study the synaptic transmission between neurons, mIPSC was recorded. 1 μM tetrodotoxin, 10 μM NBQX and 50 μM D-AP5 were added to ACSF. The same method to patch the neurons was applied and the data was recorded for 6 minutes. The amplitude and frequency were analyzed by mini-analysis software.

### RT-qPCR

All the following experiments were RNase-free. The total RNA was extracted by the traditional Trizol (Thermo, 15596018) method(Groppe and Morse 1993), dissolved in a suitable volume of RNase-free water, and the RNA concentration was determined using a Nanodrop. The RNA was then reverse-transcribed using a reverse transcription kit (Accurate Biotechnology Co., Ltd, AG11705). The cDNA solution was subjected to RT-qPCR using TB Green Premix Ex Taq (Takara, rr420a) and prime script rt master mix (Takara, rr036a). Gapdh was used as the internal control, and the relative expression of the gene was calculated by 2^−ΔΔCt^ method[62]. The primer sequences were shown in Table 1.

### mRNA sequencing and bioinformatic analyses

The original Sequencing data were analyzed using previously published methods[63, 64].The total RNA was extracted using Trizol method and the RNA integrity was determined using RNA Nano chip and Agilent 2100 bioanalyzer[65]. The samples were subjected to high-throughput sequencing through the paired-end sequencing mode of the Illumina HiSeq sequencing platform. For each sample, the pre-processing sequence was compared with the mouse genome sequence (release-98) by STAR software after removing the linker sequence fragments and low-quality fragments. For all samples, StringTie software was used to count the original sequence counts of known genes, and the expression of known genes was calculated using fragments per kilobase of transcript per million fragments mapped (FPKM). The DESeq2 software was used to screen the differentially expressed genes between different sample groups, and the differential expression range of 1 <= log_2_ (Fold Change), log_2_ (Fold Change) <= −1 and *P* value < 0.05 was used to screen the differentially expressed genes between the two groups. The function of differentially expressed genes were analyzed by David[66, 67] website (https://david.ncifcrf.gov/), including gene ontology (GO) and kyoto encyclopedia of genes and genomes (KEGG). The raw and processed data were deposited in Gene Expression Omnibus database with GEO# 157903.

### MGE cell transplantation and evaluation of migration behavior

MGEs from E13.5 wild-type embryonic brains were harvested and dissociated. The cell suspensions were incubated with AAV-scramble-GFP or AAV-shBrpf1-GFP for 2 hours at 37°C, during which the liquid was gently mixed every half hour. Each group had about 50,000 cells. The mixed liquid was centrifuged at 800 rpm for 5 minutes and then the supernatant was removed. The centrifugation step was repeated 2 to 3 times. The pellet was resuspended in 10ul DMEM supplemented with 10% FBS. The cell suspension was then injected into the neocortex of wild-type P1 host. Transplanted neurons were permitted to migrate in vivo for 35 days before analysis.

On day 35, mice were anesthetized, perfused with PBS followed by 4% PFA through the endocardium, and then the brain was carefully dissected. The brain was incubated overnight in 4% PFA, dehydrated in 30% sucrose in PBS until the sample sank to the bottom, embedded with optimum cutting temperature compound (O.C.T., Sakura,4583), and sectioned at a thickness of 20 μm. Rabbit anti-GFP (Proteintech, 1:500) and DAPI (1:1000) were used for immunofluorescence staining. The proportion of GFP^+^ cells in each layer of the cortex was analyzed after that the stained sections were photographed by confocal (leica SP8).

### Statistical Analysis

Data were analyzed statistically using ImageJ Fiji and GraphPad Prism 8.0 software. Differences between groups were tested by T test. The values of p <0.05 was considered statistically significant. *p < 0.05, **p < 0.01, and ***p < 0.001.

#### Data and reagent availability

Gene expression data are available at GEO with the accession number: GEO157903.

---

**Table S1.**
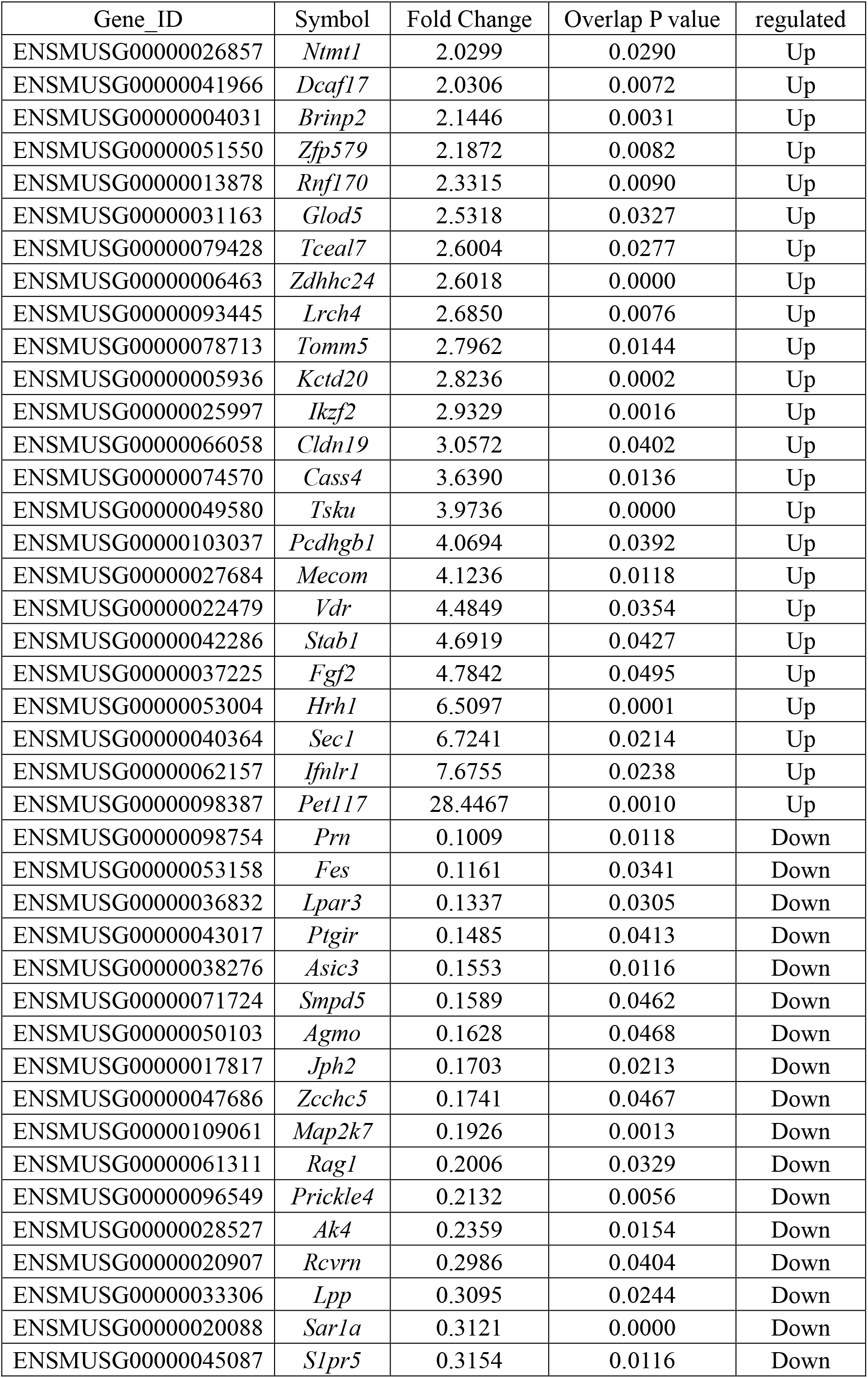
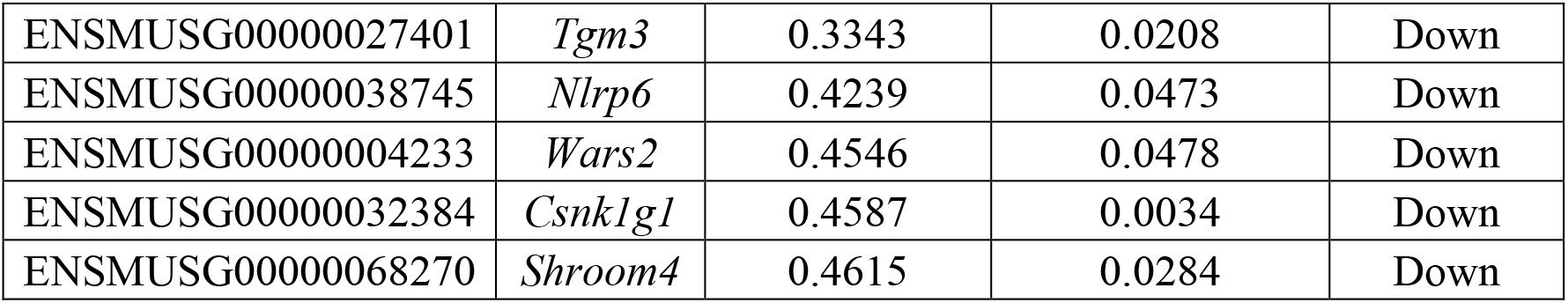
Up- and down-regulated genes by mRNA-Seq.

